# Impacts of invasive fishes on ghost frog tadpoles

**DOI:** 10.1101/2023.02.12.528228

**Authors:** Daniel van Blerk, Andrea Melotto, Josie Pegg, John Measey

## Abstract

Global amphibian populations are declining, and invasive fish are known to impact many threatened species. South Africa has both a biodiverse amphibian fauna and a rich history of invasive fish introductions with high establishment success. Studies have identified some preliminary evidence for negative impacts of invasive salmonids on two ghost frog species (Anura: Heleophrynidae). This study aims to investigate whether these negative impacts previously reported are reflected across a broader scale. Tadpoles of two species, *Heleophryne regis* and *H. purcelli* were sampled in the presence and absence of invasive fish at 112 sites across 26 streams in the Western Cape province of South Africa. A generalised linear mixed model showed invasive fish to have the most significant negative effect explaining tadpole abundance. Mean tadpole abundance decreases by 18 times in the presence of invasive fish. Environmental variables with significant effects on tadpole abundance include pH, oxygen saturation, temperature, stream depth and non-native riparian vegetation type. We can define an environmental niche of ghost frog tadpoles which does not differ between species. We conclude that invasive fish have significant negative impacts on ghost frog tadpole abundance, but the effect of the environment should not be overlooked in amphibian conservation planning and invasive species management decisions. This study supports the removal of invasive fish and alien vegetation to improve the conservation of ghost frogs.

## Introduction

A steady decline in global amphibian populations since the late 20^th^ century has led to the Amphibia being the world’s most threatened vertebrate class (Green et al., 2020). The rate of decline has been likened to a mass extinction, which could portend future losses in other vertebrate groups (Wake & Vredenburg, 2008). This decline is attributed to a broad range of current and historical causes. Current causes include climate change, pollution, and the spread of infectious disease; whilst habitat loss, commercial use and invasive species are historical causes that are important and ongoing (Collins, 2010). The impacts of historical causes are well documented but difficult to isolate as they can act in synergy, especially when invasive species are involved (Alford & Richards, 1999; Collins & Storfer, 2003; Angus, Turner & Measey, 2023). Invasive species are a concern as there is no foreseeable end to introductions of non-native species, with predictions that impacts are likely to increase in severity (Seebens et al., 2017).

Global reviews report extensive negative impacts from invasive species on native amphibians (Kats & Ferrer, 2003; Bucciarelli et al., 2014; Nunes et al., 2019; Falaschi et al., 2021). These impacts include changes to native amphibian fitness, abundance, development, growth, behaviour and morphology (Nunes et al., 2019). Mechanisms of impact include hybridisation, habitat alteration, competition and predation: the most reported interaction between invasive species and native amphibians (Bucciarelli et al., 2014; Nunes et al., 2019; Falaschi et al., 2021). The aquatic life stages of amphibians are most vulnerable to predation, especially from fish (e.g. Gillespie, 2001; Remon et al., 2016). Alien fish have a wide invasive range, a high diversity of established species and a rich history of introductions (Cambray, 2003). Impacts of introduced predators are predicted to be worse in previously fishless water bodies, as the lack of coevolutionary history with native amphibians can hamper the expression of adequate anti-predator strategies towards the novel predators (Denoel, Dzukic & Kalezic, 2005; Cox & Lima, 2006; Melotto et al., 2021). Examples of naturally fishless water bodies are headwater streams and high-altitude lakes, where fish dispersal is prevented by the steep-gradient and fast flow of water. However, predatory fish continue to be introduced to water bodies historically lacking a native counterparts (e.g. Denoel, Dzukic & Kalezic, 2005; Miró, Sabás & Ventura, 2018), resulting in the need to quantify their impacts on native amphibians.

Two study types are commonly used to determine the impacts of invasive fish on amphibians in fishless water bodies: experimental and observational studies. Experimental studies typically involve isolating the interaction between predator and prey using specially designed experiments (e.g. Shaffer & Johnson, 2008; Martín-Torrijos et al., 2016). Observational studies in the context of invasive predators impacting native amphibians may allow for a more refined investigation of the responsible impact mechanisms, and are typically surveys that draw comparisons between the native amphibian abundance, density or presence/absence in invaded and uninvaded habitats (Kats & Ferrer, 2003).

Observational studies have been used around the world to quantify invasive fish impacts on native amphibians. For example, field surveys conducted in a high-altitude Pyrenean lake attribute the exclusion of five out of six native amphibian species to the presence of invasive rainbow trout, Oncorhynchus mykiss (Miró, Sabás & Ventura, 2018). Environmental variables present a significant challenge to such assessments as the difference in amphibian abundance between a fishless and fish-invaded water body could be better explained by the habitat features instead of the presence of invasive fish. Miró et al. (2018) overcame this by sampling environmental characteristics that could influence the occurrence of amphibian species and included them in a generalised additive model to test which variables significantly affect amphibian occurrence. Similarly, Velasco et al. (2018) conducted a field study to determine the impacts of invasive fish on stream-adapted anurans in Patagonia combining habitat descriptors to invasive fish presence in their occupancy model (Velasco et al., 2018). Such studies underline the importance of collecting pertinent environmental variables and sampling a wide range of sites when comparing amphibian abundance in fish-invaded and uninvaded areas, highlighting how this approach can allow drawing much more robust conclusions from observational studies.

A review of fish invasions in South Africa found that 55 species have been introduced to the country, of which 44 have established, including predatory gamefish (Ellender & Weyl, 2014). Brown trout, *Salmo trutta* (Linnaeus 1758), and rainbow trout were introduced to South Africa for sport fishing in the late nineteenth century (Cambray, 2003). Sport fishing in South Africa remains the leading contributor to non-native fish introductions, with trout mostly being introduced into the upper reaches of mountain streams, where their impacts on resident fauna, such as native fish, have already been documented (Cambray, 2003; Shelton, Samways & Day, 2015; Khosa et al., 2019). Downstream, where warmer water and slower flow is not ideal for the persistence of trout, other gamefish species are introduced, such as smallmouth bass, *Micropterus dolomieu* (Lacépède 1802) (Ellender & Weyl, 2014). South African streams are thus under intensive stocking regimes, involving multiple predatory gamefish that are known to have had negative impacts on amphibians around the world (Cambray, 2003; Loppnow, Vascotto & Venturelli, 2013; Ellender & Weyl, 2014). Highly impactful invasive fish now occupy the same headwater streams as South African endemic anurans, such as the endemic family of ghost frogs: Heleophrynidae.

A single-site investigation of rainbow trout impacts on the Cederberg ghost frog, *Heleophryne depressa* (FitzSimons 1946), showed a decline in tadpole abundance below a waterfall where invasive rainbow trout occur (Avidon et al., 2018). Above the waterfall there was tenfold higher mean relative tadpole abundance when compared to below (Avidon et al., 2018). Similar results were found in KwaZulu-Natal, where tadpole abundance of Natal cascade frog, *Hadromophryne natalensis* (Hewitt 1913), decreased by a factor of up to 15 times at three sites where invasive brown trout occur below a physical dispersal barriers (waterfalls) (Karssing, Rivers-Moore & Slater, 2012). However, deliberate fish introductions occur above and below such barriers and therefore a more comprehensive field study, across multiple sites and streams, will determine whether the negative impacts highlighted by Avidon et al. (2018) and Karssing et al. (2012) are more pervasive among ghost frogs. In this study, we aim to conduct such a study determining the impacts of invasive fishes on two heleophrynid species assessing tadpole abundance across multiple sites in the presence and absence of invasive fish, while accounting for environmental variables that could affect the observed tadpole abundance.

## Materials and Methods

### Study species

Ghost frogs (Anura: Heleophrynidae) contain seven species in two genera (*Heleophrynus* and *Hadromophryne*) endemic to South Africa, where they inhabit the fast-flowing headwaters of streams and rivers in fynbos and forest biomes (Minter et al., 2004; Conradie & Conradie, 2015). The tadpoles have a dorso-laterally flattened body to reduce drag in the current, and a large oral sucker is used to cling to the rocky substrate and to feed on algal film (Lukas, 2021). Tadpoles spend between 18 and 24 months in the larval stage, requiring permanent flow to survive. This permanence of flow, along with a mean annual water temperature that does not exceed 17.2°C were identified as key habitat descriptors for tadpoles of the Table Mountain ghost frog, *Heleophryne rosei* (Hewitt 1925) (Ebrahim, de Villiers & Measey, 2020). Given their specialist life history and limited suitable habitat, ghost frogs are susceptible to population declines hrough either human mediated threats (invasive species, habitat loss) or natural phenomena (floods, fires) (SAFRoG & IUCN-ASG, 2016a,b). These threats have led to *H. rosei* and Hewitt’s ghost frog, *H. hewitti* (Boycott 1988) being listed by the IUCN as Critically Endangered and Endangered, respectively (SAFRoG & IUCN-ASG, 2016a,b). The current study sampled tadpoles listed by the IUCN as Least Concern: the royal ghost frog *Heleophryne regis* (Hewitt 1909) and the Cape ghost frog *Heleophryne purcelli* (Sclater 1898).

Highly impactful invasive salmonids (Family: Salmonidae) and black bass (Family: Centrarchidae), introduced as gamefish, occupy many South African streams (Ellender & Weyl, 2014), and are known to have significant negative impacts on native amphibian larvae, macroinvertebrates and endemic fish species (Woodford & Impson, 2004; Woodford et al., 2005a; Weyl et al., 2010; Ellender, 2013; Rivers-Moore, Fowles & Karssing, 2013; Avidon et al., 2018). The documented negative impacts of invasive bass on endemic fish served as the basis for chemical and mechanical eradication from small headwater streams in the Cape Floristic Region (Weyl et al., 2014a; van der Walt et al., 2019), and the subsequent increase in endemic fish and invertebrate biodiversity (Weyl et al., 2014b; van der Walt et al., 2019; Bellingan et al., 2019). A similar population recovery in native fish followed the mechanical eradication of spotted bass from the Thee River (van der Walt et al., 2019). However, no fish removal has yet been conducted in streams with threatened amphibian species in South Africa.

### Study area

The Cape Fold Mountains are a continuous chain of quartzitic sandstone massifs that lie between the southeastern coast of Africa and the Karoo basin. From December 2021 to May 2022, we sampled 112 sites across 26 streams in an east-west section of the Cape Fold Mountains within the distributions of *H. regis* and *H. purcelli* (Figure 1). Streams sampled generally ran north-south and rose on average 345 m asl. Thirteen streams had invasive fish present at 18 sites. Native fish were present at 20 sites and 78 sites contained no fish.

**Figure 1:**
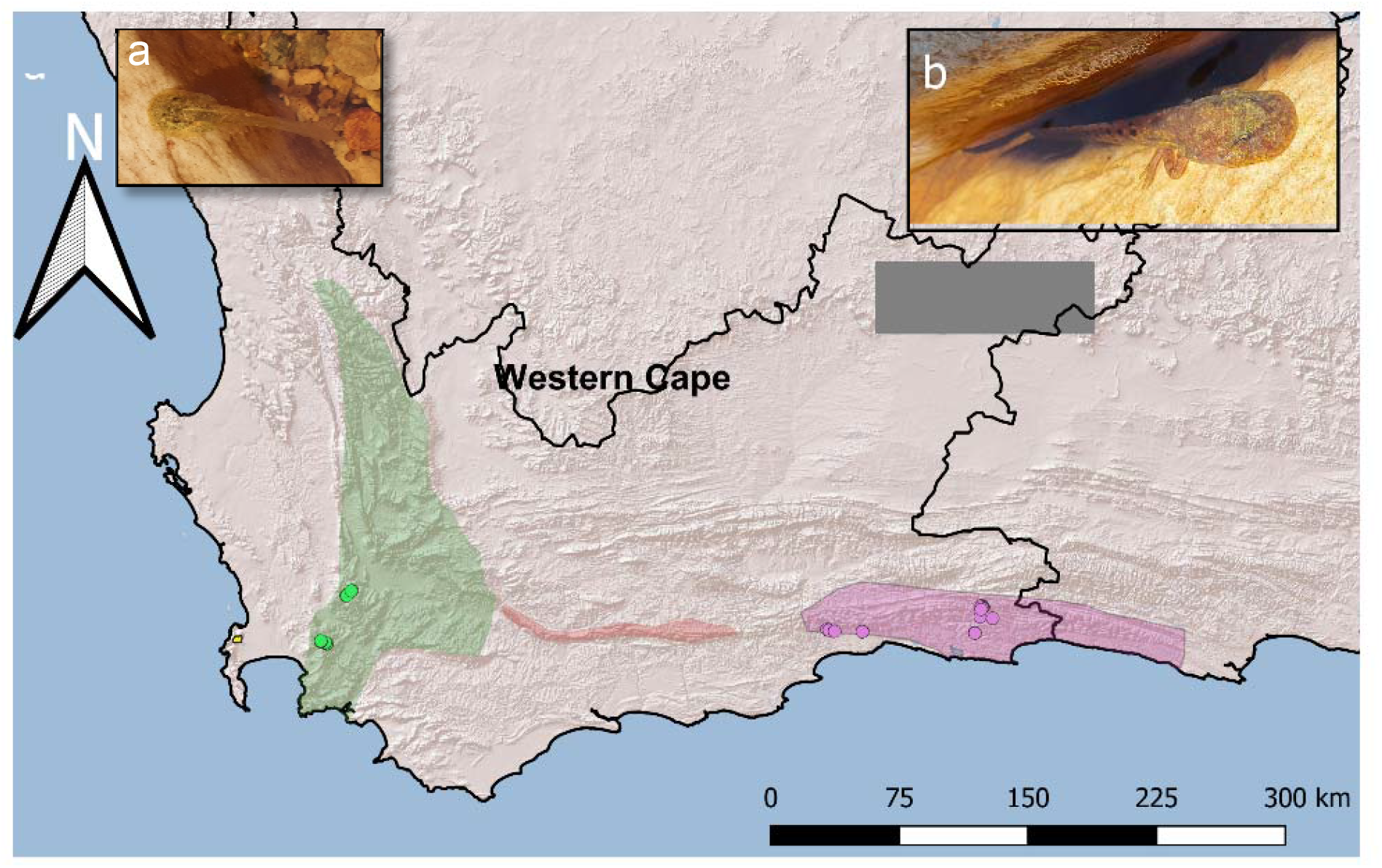
(a) Ghost frog tadpole sampling sites denoted by green (*Heleophryne purcelli*) and purple (*H. regis*) points. Polygons represent the distribution range of ghost frog species *H. rosei* (yellow), *H. purcelli* (green), *H. orientalis* (red) and *H. regis* (purple). (b) *H. regis* tadpole.

Uninvaded streams were selected based on compiled heleophrynid occurrence data obtained from the Centre of Excellence for Invasion Biology (CIB), iNaturalist and the Animal Demography Unit (ADU). Invaded streams were selected based on where the heleophrynid occurrence data overlapped with distributional data for *O. mykiss* and *M. dolomieu*, obtained from the Freshwater Biodiversity Information System (FBIS). Explorative trips were conducted to the identified streams to delineate sample sites before data was collected. We moved from downstream to upstream sites, marking the furthest point travelled upstream as the last site to be sampled. The furthest point upstream is the point where there was no more water, or a physical barrier prevented further travel upstream. The lower end sampling point was marked as the point where the headwater stream joined its larger tributary. Possibly due to the time of year, most streams did not reach a tributary and instead reached a dry point. In this case, the dry point was taken as the lowest point of the headwater stream. Along each stream, adjacent sample sites were set approximately 20 metres apart. Moving upstream ensured no interference with the water quality readings, whilst the distance between sites ensured no upstream biota would be disturbed by sampling of the downstream site.

### Tadpole Sampling

Ten-minute time limited searches were conducted at each site to sample tadpoles. The sole sampler (DvB) moved across the full width of the stream, wiping under loose stones, rocks, and boulders to wash tadpoles into a handheld net (net frame dimensions: 56 × 44 cm, mesh size: 4 mm) downstream. The area of the stream searched was noted.

### Fish Sampling

At each site, ten-minutes of electrofishing was conducted to sample fish using a SAMUS Electrofisher 725MP, using pulsed DC current at a voltage adjusted based on stream conductivity. Stunned fish were caught using a hand net and were transferred to a tray of aerated water, identified to species level, photographed and released.

### Water quality variables

A calibrated Aquaread Aquaprobe AP-5000 was placed in the middle of stream, allowing for deep enough water for the measuring chamber to fill at each site to measure the following variables: water pH (to the nearest 0.1 pH), water temperature (to the nearest 0.1 °C), electroconductivity (EC: to the nearest µS.cm^-1^), Oxygen saturation (to the nearest 0.1 %), and total dissolved solids (TDS) (to the nearest mg.L^-1^). Each measurement was stabilised for at least 5 seconds before recording. Stream surface velocity was measured using a float that was released on the upstream side of a meter rule and the time it took to reach the downstream end was recorded with a stopwatch. The average of the three measurements in the middle and either side of the stream was converted to meters per second.

### Physical site features

The mean width of the stream at each site was measured (to the nearest 0.1 m) using a measuring tape at the start, middle and end of where tadpole sampling occurred. Mean depth of the stream at the middle of each width measurement (to the nearest 0.01 m) was made with a measuring tape. Substrate type was recorded by applying size clast standards according to Rowntree and Wadeson (2000) into one of the following categories: Very Fine Sand/Silt, Fine/Medium Sand, Coarse/Very Coarse Sand, Very Fine/Fine Gravel, Medium Gravel, Coarse/Very Coarse Gravel, Small Cobble, Large Cobble, Small Boulder, Medium Boulder, Large/Very Large Boulder, Bedrock. The surface area of each site was measured in m^2^ by multiplying the width of the stream the distance travelled upstream during the manual tadpole sampling. The dominant riparian vegetation type at each sample site was assigned to one of three categories: Indigenous Forest, Fynbos or Plantation. Plantations consist of alien pines (*Pinus* spp.) or eucalypts (*Eucalyptus* spp.) trees that are often invasive (Richardson et al., 2007). Fynbos is a mediterreanan scrub biome endemic to the southwestern Cape of South Africa (see Mucina & Rutherford, 2006). Indigenous forest grows in many steep valleys within the Fynbos and also in the forest biome, and is dominated by *Podocarpus* spp. The altitude of each site was recorded in metres above sea level (asl) with the GPS function of the Aquaread Aquaprobe AP-5000.

### Statistical Analysis

#### Independence of habitat variables: Correlations

Correlation tests were conducted to determine which of the sampled habitat variables were strongly correlated (r > 0.3) and should be considered for exclusion of the analysis. The tests were conducted using the *psych* package (Revelle 2022) in R (R Core Team, 2021). Correlation tests showed a strong correlation (r > 0.5, p < 0.05) of total dissolved solids (TDS) with pH and further significant correlations (p < 0.05) with temperature, flow and altitude (Table S1). Accordingly, TDS was removed from further analysis and pH retained because water acidity has been associated with effects on performance, growth and survivorship in other stream breeding amphibian larvae (Clark & Lazerte, 1985; Green, Thompson & Lemckert, 2004). Other relationships that showed some degree of correlation (p < 0.05) include pH-Temperature, pH-Oxygen Saturation, pH-Altitude, Temperature-Electroconductivity, Temperature-Width, Temperature-Altitude, Flow-Depth, Width-Depth, Width-Area, Depth-Area (Table S1). However, none of the variables in these relationships were excluded as they were not strongly correlated (r < 0.3).

#### Independence of environmental variables: Collinearity

Variance Inflation Factor (VIF) were used to assess potential multicollinearity among the collected environmental predictor variables. VIF is an index estimating how much the variance of regression coefficients is affected by collinearity (Craney & Surles, 2002). For each predictor included in the regression, an estimated VIF value provides its relative contribution to collinearity given the simultaneous presence of the other ones (Zuur, Ieno & Elphick, 2010). Although there is a lack of consensus in the literature on when a VIF value is significant, a VIF threshold value of <3 is generally considered to cause negligible collinearity effects, and we adopted this threshold to assess potential collinearity among predictors of this study (Craney & Surles, 2002; Zuur, Ieno & Elphick, 2010; Akinwande, Dikko & Samson, 2015). VIF values were obtained using the *car* package (Fox & Weisberg, 2019). Potential collinearity was inspected following (Zuur, Ieno & Elphick, 2010). VIF values were initially estimated including all environmental predictors. Then, the predictor with the highest value among those ones showing a VIF > 3 was discarded, and VIF was calculated again for the remnant ones. This process was repeated until no predictors with VIF > 3 were present, and this subset of predictors was used in the subsequent analyses to assess the environmental features affecting tadpole abundance.

This procedure showed plant coverage had a significant VIF value with both pH and temperature (VIF > 3). Consequently, plant coverage was removed from the model, as temperature is related to overstory cover (Moore, Spittlehouse & Story, 2005), while we kept temperature as its effects on amphibian ontogenesis are widely documented (e.g., Wassersug, 2000), and it is reported to have a significant influence on development and condition of heleophrynid larvae (Rivers-Moore, Fowles & Karssing, 2013).

#### Independence of species

To ensure that there was no significant difference between *H. regis* and *H. purcelli* in the habitat and environmental data, we fitted a generalised linear mixed model with binomial distribution to the response variable of frog species, using the *glmmTMB* package (Brooks et al., 2017) in R.

The binomial species model identified only altitude as having a significant effect on ghost frog species due to the five highest sites only occurring within the range of *H. purcelli*. The range of mountains containing *H. regis* does not obtain the altitude commensurate with those of *H. purcelli* (see Figure 1). On removal of these five highest sites from the data set, no differences in expected mean value of dependent variables was found, and so both species of ghost frog tadpoles were combined in the model. It is worth noting that these five highest sites were from the same stream (lower sites were retained) and had only tadpoles and no invasive fish recorded.

#### The effect of invasive fish and environmental variables on tadpole abundance: Direction and significance

To determine the effect of the sampled habitat variables and invasive fish on ghost frog tadpoles, we fitted generalised linear mixed effect models (GLMMs) with negative binomial zero-inflated distributions (Brooks et al., 2017) with tadpole abundance as response variable. Fixed effects included in the model were pH, temperature, electroconductivity, oxygen saturation, average flow speed, clast size (continuous, ranked number assigned to corresponding substrate type in order of smallest to largest), area, average depth, average width, altitude, vegetation type (i.e., indigenous forest, fynbos, plantation) and invasive fish presence. The stream in which the site occurs was included as a random factor. Relative support for the model was assessed through the calculation of Akaike’s information criterion (AIC). AIC values were compared between models varying in complexity (number of fixed effects), all being fitted to the same variable of interest (tadpole abundance). The models were built with the *glmmTMB* package (Brooks et al., 2017) in R.

Mean tadpole abundance at fish invaded sites were compared to uninvaded sites to determine effect size. This method of reporting effect size was used in previous literature that quantified impacts of invasive fish on ghost frog tadpoles (Karssing, Rivers-Moore & Slater, 2012; Rivers-Moore, Fowles & Karssing, 2013; Avidon et al., 2018).

## Results

The model with most fixed effects was chosen as the final tadpole abundance model based on the lowest AIC value (see Table S2). The results of this model show that the presence of invasive fish had a significant negative impact on ghost frog tadpole abundance (Figure 2). Water quality variables with significant effects on tadpole abundance include pH, Temperature and Oxygen Saturation (Figure 2). No tadpoles occurred above 19°C, with the highest tadpole abundances found between 16°C and 19°C (Figure S1). No tadpoles were found above a pH of 6.9, with the most tadpoles found between pH of 3 and 5.5 (Figure S2). This implies that more acidic streams are associated with higher tadpole abundance. Higher tadpole abundance is associated with an increase in oxygen saturation (Figure 2). Depth is the only physical stream characteristic to have a significant effect on tadpole abundance, with a decrease in tadpole abundance associated with an increase in depth (range: 4.33cm to 226.33cm) (Figure 2). To better visualise the effect of vegetation type, post-hoc two-way comparisons between Fynbos and Indigenous Forest, Fynbos and Plantation, and Indigenous Forest and Plantation were added to Figure 2. The two-way comparison between vegetation types shows that there are significantly more tadpoles in fynbos and indigenous forest when separately compared to alien plantations. Although there are more tadpoles in fynbos than indigenous forest, the difference is not significant (Figure 2).

**Figure 2:**
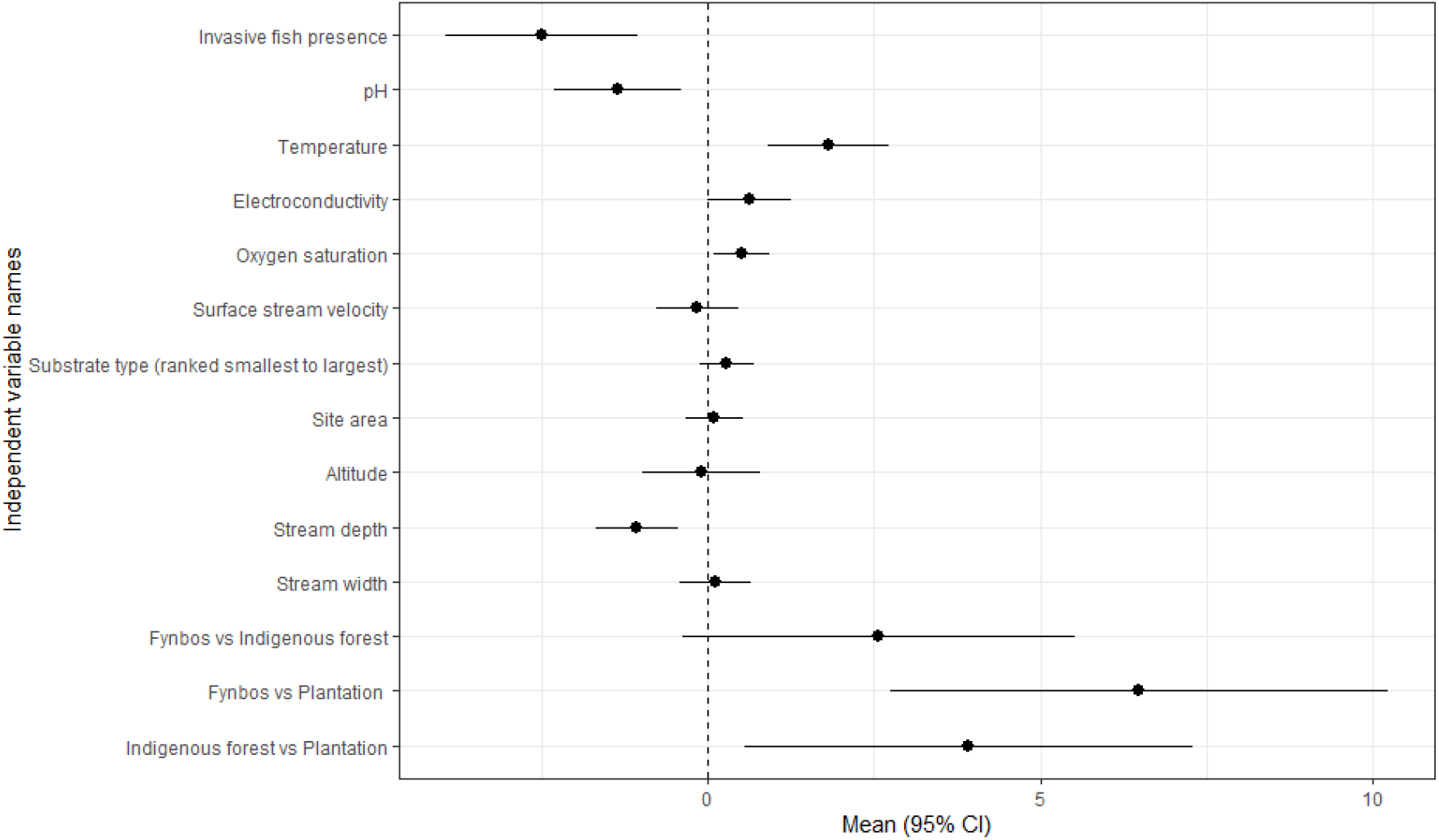
Forest plot depicting the significance and direction of the model independent variables’ effect on ghost frog (Heleophrynidae) tadpole abundance. Independent variable names are listed on the y-axis, with the last three lines representing two-way comparisons between the categories of riparian vegetation types recorded in the field. Mean model intercepts are on the x-axis. The points in line with the independent variable represents the mean intercept with 95% confidence interval. The dashed vertical line represents the line of null effect. An effect to the right of the line of null effect has a positive effect on tadpole abundance. Negative effects on tadpole abundance will have a mean intercept to the left of the line of null effect. An effect is considered significant when the confidence intervals do not cross the line of null effect.

Mean tadpole abundance decreased by ∼18 times in the presence of invasive fish, whilst tadpole density decreased by ∼4 times when invasive fish were present (Table 1).

**Table 1:** Mean ghost frog tadpole abundance (± Standard Error) and density (per m^2^) at sites in the southern Cape Fold Mountains where invasive fish are present and absent.

|  | Mean tadpole abundance | Mean tadpole density |
| --- | --- | --- |
| Invasive fish absent | 7.00 ±1.603 | 2.00 ±0.537 |
| Invasive fish present | 0.389 ±0.2161 | 0.072 ±0.0394 |

## Discussion

Field sampling across 112 sites within the distribution range of two heleophrynids that we conducted for this study supports the hypothesis that invasive fish have significant negative impacts on ghost frog tadpole abundance, and adds to evidence of negative impacts of invasive fish on amphibians globally (Bucciarelli et al., 2014; Nunes et al., 2019; Falaschi et al., 2020). The results corroborate previous studies on ghost frog tadpoles that reported negative impacts of invasive salmonids at a smaller scale (Karssing, Rivers-Moore & Slater, 2012; Avidon et al., 2018). Here we report negative effects of invasive fish on *H. purcelli* and *H. regis*, both listed by the IUCN as Least Concern, but similar impacts likely apply to all heleophrynid species, including the Critically Endangered and Endangered species, *H. rosei* and *H. hewitti*. Moreover, given the extent of fish invasions within the distribution ranges of *H. purcelli* and *H. regis*, a reassessment of their conservation status is now urgently needed. This puts an imperative on collecting accurate records on the presence of invasive fish (for example, using eDNA:) throughout the range of this South African endemic family.

Additionally, we provide valuable information on what constitutes suitable habitat for heleophrynid tadpoles. Acidic streams are associated with higher tadpole abundance, as previously suggested (Ebrahim, de Villiers & Measey, 2020). The pH of a stream is affected by terrestrial biotic mechanisms, including acidification of water by riparian vegetation such as in the fynbos biome, with indigenous forest and pine plantations also having the capacity to acidify water (Midgley & Schafer, 1992).

However, this relationship between water pH and tadpole abundance may also reflect the preferences of adult ghost frogs for the surrounding terrestrial environment, and not necessarily a physiological requirement of tadpoles. This is supported by the higher abundance of tadpoles observed in fynbos and indigenous forest dominated landscapes found in this study. Another biotic mechanism responsible for increased tadpole abundance in sites with lower pH would favour acid tolerant algal species and indirectly increase the environmental suitability for organisms grazing upon them, such as ghost frog tadpoles. Alternatively, the abiotic mechanism through which tadpole abundance increases in acidic streams could depend on the inverse relationship of pH with temperature and dissolved oxygen (Moore, Spittlehouse & Story, 2005), supported in line with the significant positive effect of temperature and oxygen saturation reported in the present study. The biotic and abiotic mechanisms through which pH can change are not mutually exclusive, and the sole importance of pH for heleophrynid tadpoles remains to be tested, for instance by investigating their survival and performance in controlled laboratory conditions.

The positive effect of water temperature is unexpected, as previous literature reports 17.2°C to be the upper limit for tadpoles of a congeneric species, such as *H. rosei* (Ebrahim, de Villiers & Measey, 2020). It is, however, important to consider the range of temperatures sampled in this study is 10.1°C – 23.9°C (Figure S1). It is possible that the 19°C threshold in the current study is due to thermal tolerance differences between ghost frog species. Research on the Natal cascade frog reports even higher annual water temperatures (20+°C) for sample sites in which tadpoles persist, suggesting that these tadpoles may be more resilient to warm temperatures than reported (Rivers-Moore & Karssing, 2014). The negative effect of increasing depth on tadpole abundance could be due to presence of invasive fish predators, which instead are absent or rarer in shallow waters. This is supported by the larger proportion of invasive fish presence at the deeper sampled sites (Figure S3). The significant effect of riparian vegetation type on tadpole abundance shows that alien pine plantations are associated with a decrease in tadpole abundance, either because of a direct effect on the aquatic habitat or because they alter the terrestrial habitat for adults. Ebrahim et al. (2020) hypothesise that pine plantations slow the flow of water and provide less shading to the stream, which negatively affects tadpole occurrence of *H. rosei*. This effect specifically on heleophrynid tadpoles remains untested, but pine and eucalyptus plantations have shown to significantly reduce water flow in South African streams during the first 6 to 20 years of tree growth (Scott & Prinsloo, 2008). However, once plantation trees reach 40 years of age, water flow is not significantly different to pre-afforestation measurements (Scott & Prinsloo, 2008). This could explain the persistence of *H. hewitti*, which is exclusively found in four plantation streams in the Eastern Cape and has exhibited population stability (Conradie & Conradie, 2015). Since there is no natural vegetation left for *H. hewitti*, experimental plots cleared of alien vegetation are being used to investigate the effects of the plantations on tadpoles (Conradie & Conradie, 2015). The results of the present study suggest that the densities of *H. hewitti* tadpoles might increase if pine plantations were removed from the vicinity of streams.

Although the current study provides conclusive evidence for the impact of invasive fish, there are several caveats that should be considered. One limitation to the study is the seasonal nature of the sampling. Monitoring tadpole abundance in the presence of invasive fish throughout the year would provide a broader picture and could determine whether there is any seasonal nature to the impact of invasive fish on heleophrynid tadpoles. However, the field season for headwater stream sampling is restricted to summer because of difficulties related to decreased tadpole detectability and danger to personnel associated with the winter rains. Seasonal monitoring using area counts of *H. rosei* tadpoles has been conducted each summer since the 1990s for this reason (Measey, 2011; Measey et al., 2019). Yearlong monitoring of heleophrynid tadpole populations has been conducted on *H. natalensis* and *H. hewitti* (Rivers-Moore & Karssing, 2014; Conradie & Conradie, 2015), but not with the aim of recording invasive fish impacts. Year-round monitoring programmes for other heleophrynid species are needed, and we also recommend contextual recording of invasive fish data to investigate potential seasonal dependency of their impacts.

Another limitation in the sampling conducted for this study is the lack of data on native predators (e.g., dragonfly nymphs, freshwater crabs, *Anguilla* spp.) that are likely to prey upon ghost frog tadpoles and possibly contribute to explain differences in their abundance among sites. Three native fishes were found during the current study (see Table S3), including two redfin species: the Eastern Cape redfin, *Pseudobarbus afer* (Peter 1864) and the slender redfin *P. tenuis* (Barnard 1938). No tadpoles were found in association with the two redfin species, but this may be due to the high pH, temperature, and depth of the sites and not due to exclusion via redfin predation. This prevented analysis on the influence of native fish. The only other native fish species found co-occurring with ghost frog tadpoles in this study was *Galaxia zebratus* (Castelnau 1861); from a genus of small bodied insectivorous fish that show high species endemism to headwater streams in the Western Cape (Chakona, Swartz & Gouws, 2013; Ellender et al., 2017). Severe declines in population and total exclusion of species from the genus *Galaxias* are associated with rainbow trout and smallmouth bass invasions in South Africa (Woodford & Impson, 2004; Shelton, Samways & Day, 2015). Research on the native fish predators of heleophrynid larvae would contribute towards understanding factors potentially influencing tadpole distribution and abundance. The main mechanism through which invasive fish negatively impact on ghost frog tadpole abundance is suspected to be predation with direct removal of larval stages and consequent recruitment failure, but this has not been directly tested in this study or elsewhere, and other mechanisms may exist. For instance, an alternative hypothesis is that the presence of invasive predatory fish influences breeding site choice of adult ghost frogs, leading to lowered tadpole abundance or absence.

Our study will help conservation managers make decisions to protect amphibians. A review of conservation decisions reports that primary scientific literature is the least cited source for conservation actions (Sutherland et al., 2004). Most decisions are made on anecdotes without consulting primary literature, which is a high-risk strategy for threatened endemic species. The present study alone is not enough to for multilateral action in the evidence-based approached proposed by Sutherland et al. (2004). The effect of the environment and habitat on ghost frog tadpole abundances reported in this study shows that not all streams invaded by predatory fish are suitable for ghost frog tadpoles. To bridge the gap between current knowledge and conservation decision making, future research is needed on a catchment-by-catchment basis. The lack of data on invasive fish distributions was apparent when selecting sites for field sampling, and such information represents a precursor to future management decisions.

## Conclusions

Our data show that invasive fish and other environmental factors (temperature, pH, depth, and pine plantations) have significant impacts on ghost frog tadpole abundance. The scale of this study suggests that the negative effect of invasive fish on ghost frog tadpoles is likely reflected across all ghost frog species, including the IUCN threatened species *H. rosei* and *H. hewitti*. Our results are sufficient to require the reassessment of the impact of alien fish on these and other heleophrynids, but this will require comprehensive data on the presence of invasive fish within their ranges (e.g., through eDNA). Catchment specific studies on species under threat would be required to justify management interventions on the scale of invasive fish removal (Woodford et al., 2005b; van der Walt et al., 2019). This study represents a step towards evidence-based conservation as proposed by Sutherland et al. (2004). The effect of water quality and habitat descriptors on ghost frog tadpole abundance reported in this study need to be assessed without the influence of invasive fish. More research is required on the mechanisms involved in the impact of invasive fish on heleophrynids to bridge the gap between current knowledge and promote evidence-based conservation decision-making.

## Acknowledgements

We would like to thank the many landowners and clubs that provided information or gave their assent for us to sample on their land. We dedicate this work to the memory of Prof Olaf Weyl for his inspirational work on invasive fish in South Africa and his initial enthusiasm for this project.

## Supplementary information

**Table S1:** r value and p values for independent variables included in the analysis between tadpole abundance and environmental variables of sites visited during the study.

|  | ph | temp | cond | oxsat | tds | flowavera | widthaver | depthaver | clastsized | area | altitude |
| --- | --- | --- | --- | --- | --- | --- | --- | --- | --- | --- | --- |
|  | r | r | r | r | r | r | r | r | r | r | r |
| ph | 1 | 0.263597 | -0.05782 | 0.39007 | -0.60368 | 0.200789 | -0.00158 | 0.138054 | -0.08069 | 0.089257 | 0.251512 |
| temp | 0.263597 | 1 | 0.569537 | 0.190834 | -0.20836 | -0.08636 | 0.268392 | -0.02039 | 0.053033 | 0.150855 | -0.36671 |
| cond | -0.05782 | 0.569537 | 1 | -0.00971 | -0.032 | -0.05061 | 0.117707 | -0.02947 | 0.084436 | 0.029115 | -0.04783 |
| oxsat | 0.39007 | 0.190834 | -0.00971 | 1 | -0.20513 | -0.17896 | -0.06924 | -0.10425 | 0.068265 | -0.14375 | 0.085114 |
| tds | -0.60368 | -0.20836 | -0.032 | -0.20513 | 1 | -0.22554 | 0.120487 | -0.03529 | 0.099338 | -0.08642 | -0.29787 |
| flowavera | 0.200789 | -0.08636 | -0.05061 | -0.17896 | -0.22554 | 1 | -0.05423 | 0.191421 | 0.050234 | 0.000765 | 0.048535 |
| widthaver | -0.00158 | 0.268392 | 0.117707 | -0.06924 | 0.120487 | -0.05423 | 1 | 0.31819 | 0.127397 | 0.703676 | -0.11487 |
| depthaver | 0.138054 | -0.02039 | -0.02947 | -0.10425 | -0.03529 | 0.191421 | 0.31819 | 1 | 0.122866 | 0.247749 | 0.007787 |
| clastsized | -0.08069 | 0.053033 | 0.084436 | 0.068265 | 0.099338 | 0.050234 | 0.127397 | 0.122866 | 1 | 0.068176 | 0.028578 |
| area | 0.089257 | 0.150855 | 0.029115 | -0.14375 | -0.08642 | 0.000765 | 0.703676 | 0.247749 | 0.068176 | 1 | 0.077355 |
| altitude | 0.251512 | -0.36671 | -0.04783 | 0.085114 | -0.29787 | 0.048535 | -0.11487 | 0.007787 | 0.028578 | 0.077355 | 1 |
|  | pval | pval | pval | pval | pval | pval | pval | pval | pval | pval | pval |
| ph | 0 | 0.29751 | 1 | 0.001859 | 4.00E-10 | 1 | 1 | 1 | 1 | 1 | 0.427929 |
| temp | 0.00633 | 0 | 9.95E-09 | 1 | 1 | 1 | 0.259531 | 1 | 1 | 1 | 0.005635 |
| cond | 0.556012 | 1.88E-10 | 0 | 1 | 1 | 1 | 1 | 1 | 1 | 1 | 1 |
| oxsat | 3.57E-05 | 0.050053 | 0.921312 | 0 | 1 | 1 | 1 | 1 | 1 | 1 | 1 |
| tds | 7.40E-12 | 0.032083 | 0.744718 | 0.034913 | 0 | 0.884078 | 1 | 1 | 1 | 1 | 0.09448 |
| flowavera | 0.039037 | 0.378736 | 0.606407 | 0.066432 | 0.020093 | 0 | 1 | 1 | 1 | 1 | 1 |
| widthaver | 0.987142 | 0.005407 | 0.229489 | 0.480616 | 0.218598 | 0.58084 | 0 | 0.044376 | 1 | 2.21E-15 | 1 |
| depthaver | 0.158165 | 0.835682 | 0.76428 | 0.28754 | 0.719512 | 0.049339 | 0.000888 | 0 | 1 | 0.470292 | 1 |
| clastsized | 0.410962 | 0.589256 | 0.389485 | 0.486863 | 0.310999 | 0.609083 | 0.193123 | 0.209571 | 0 | 1 | 1 |
| area | 0.362888 | 0.12269 | 0.76703 | 0.141545 | 0.378373 | 0.993787 | 4.03E-17 | 0.010451 | 0.48743 | 0 | 1 |
| altitude | 0.009303 | 0.00011 | 0.626339 | 0.385673 | 0.001928 | 0.62126 | 0.241002 | 0.936859 | 0.771205 | 0.430606 | 0 |

**Table S2:**
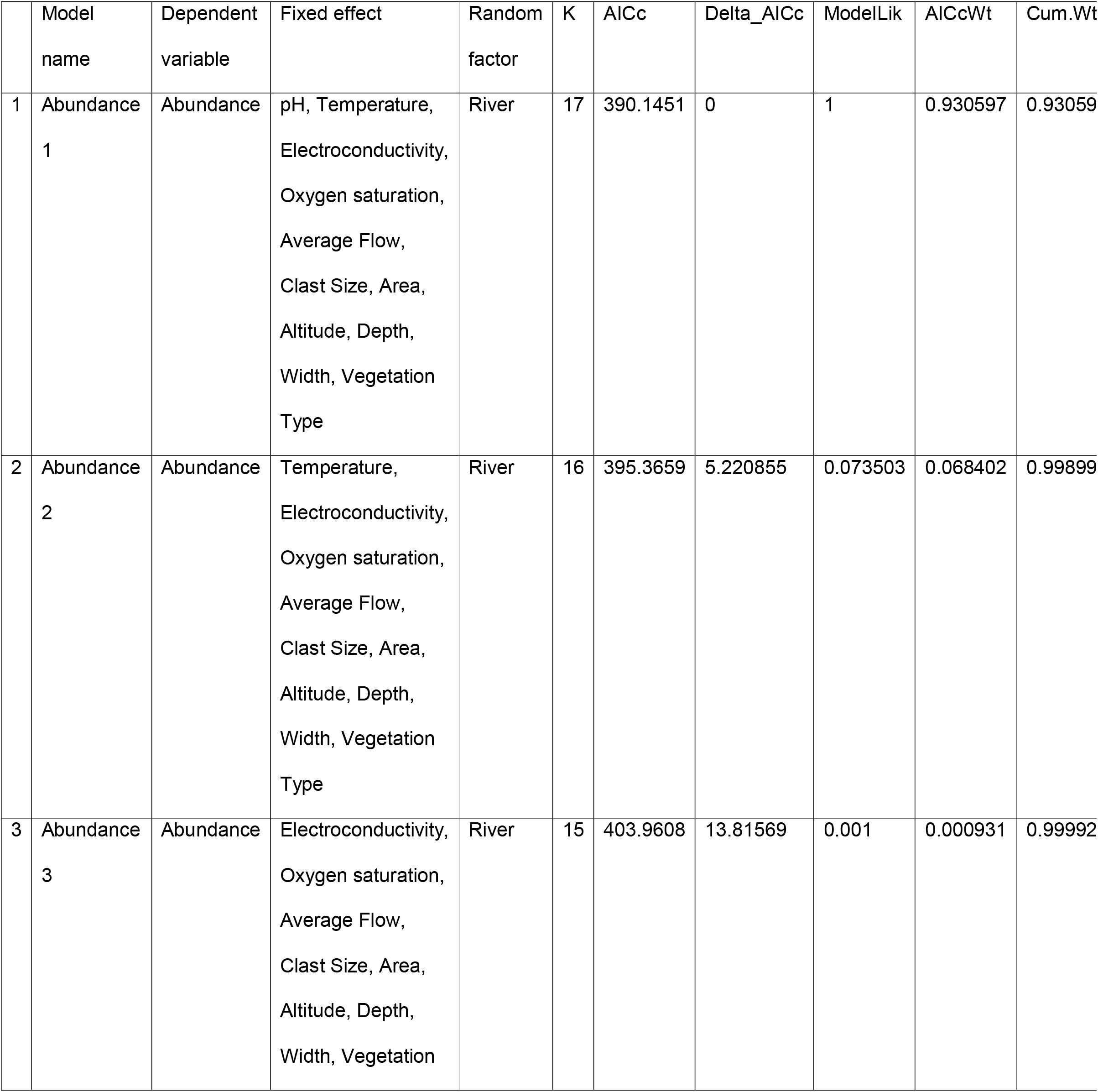
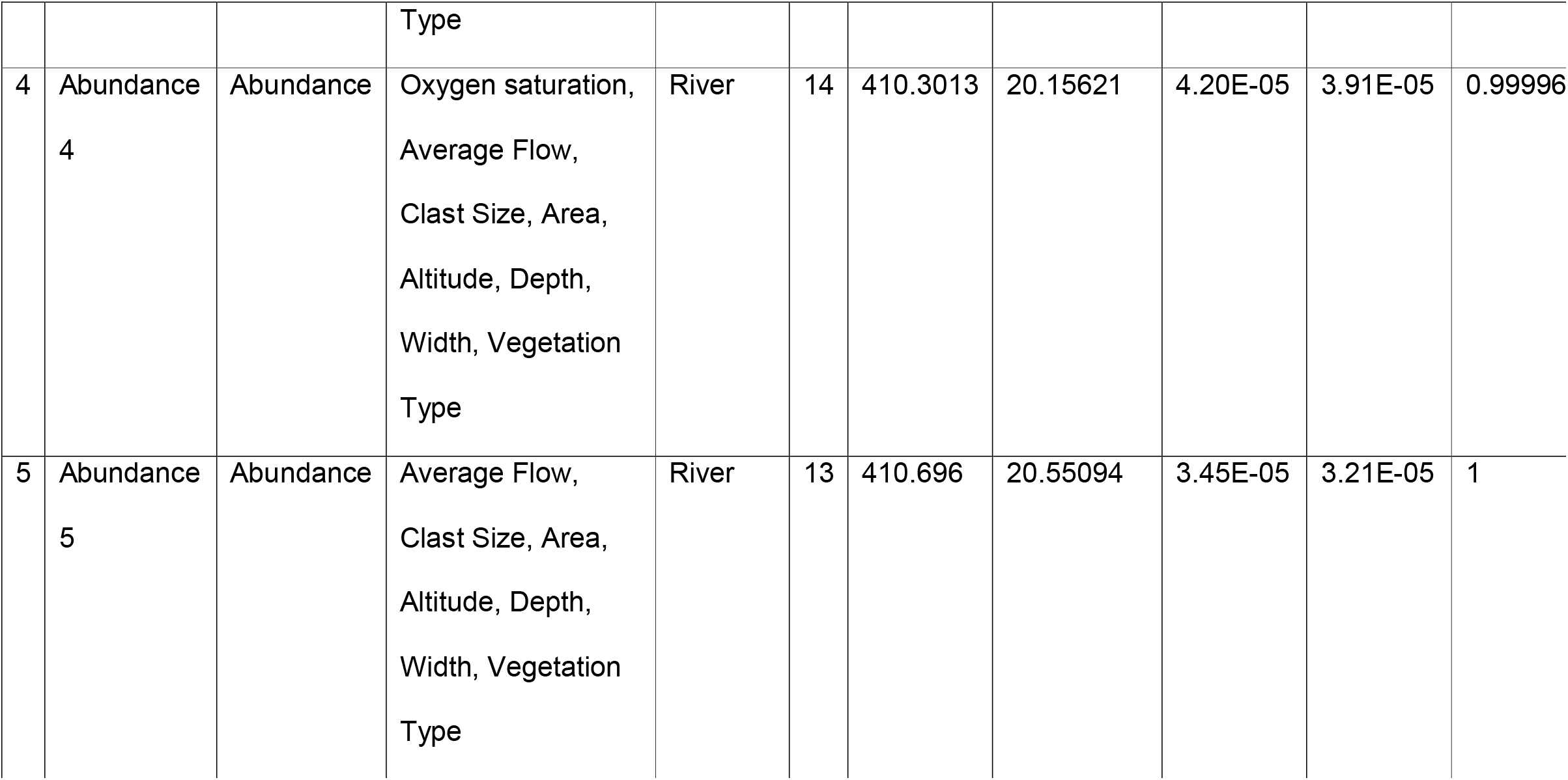
Summary of tadpole abundance models. For each model, Model number and name, Dependent variable, Fixed effect, Random factor, Number of estimated parameters (K), Akaike’s Information Criterion (AICc), Delta AIC, Relative likelihood of model (ModelLik), Akaike weights (AICcWt), Cumulative Akaike Weights.

**Table S3:** List of fish species recorded during this study sampling ghost frog tadpoles in the southern Cape Fold Mountains. The status denotes whether each species is within its native or invasive range.

| Family | Species | Common name | Status |
| --- | --- | --- | --- |
| Salmonidae | *Oncorhynchus mykiss* | Rainbow trout | Invasive |
| Salmonidae | *Salmo trutta* | Brown trout | Invasive |
| Centrarchidae | *Micropterus spp.* | Black bass | Invasive |
| Cichlidae | *Tilapia sparrmanii* | Banded tilapia | Invasive |
| Galaxiidae | *Galaxia zebratus* | Cape zebra | Native |
| Cyprinidae | *Pseudobarbus afer* | Eastern cape redfin | Native |
| Cyprinidae | *Pseudobarbus tenuis* | Slender redfin | Native |

**Figure S1:**
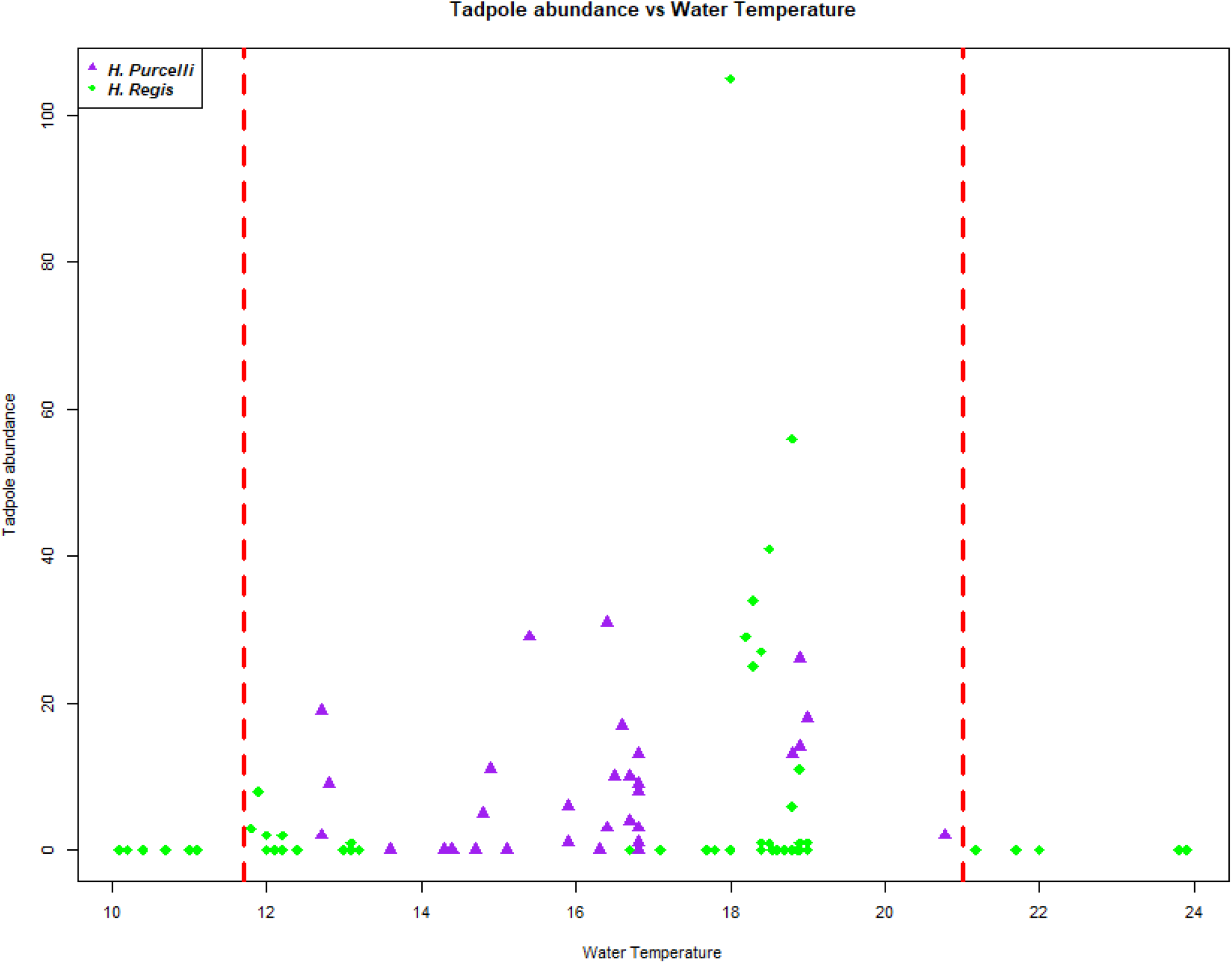
Scatterplot of ghost frog tadpole abundance against temperature at each site sampled in the southern Cape Fold Mountains. Red vertical dashed lines represent the range in which at least 1 tadpole was found.

**Figure S2:**
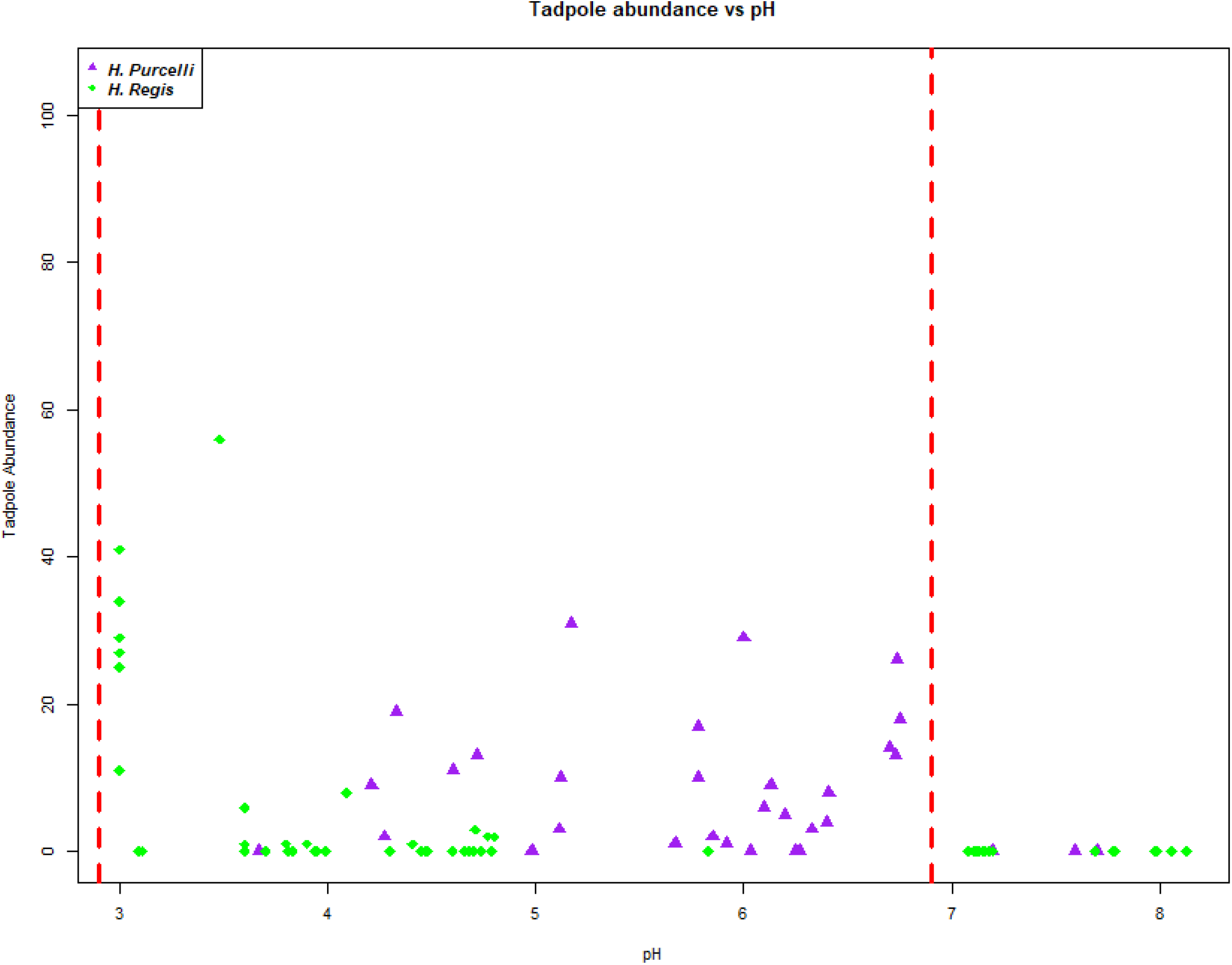
Scatterplot of ghost frog tadpole abundance (see methods for interpretation of units) against pH at each site. Red vertical dashed lines represent the range in which at least 1 tadpole was found in the southern Cape Fold Mountains.

**Figure S3:**
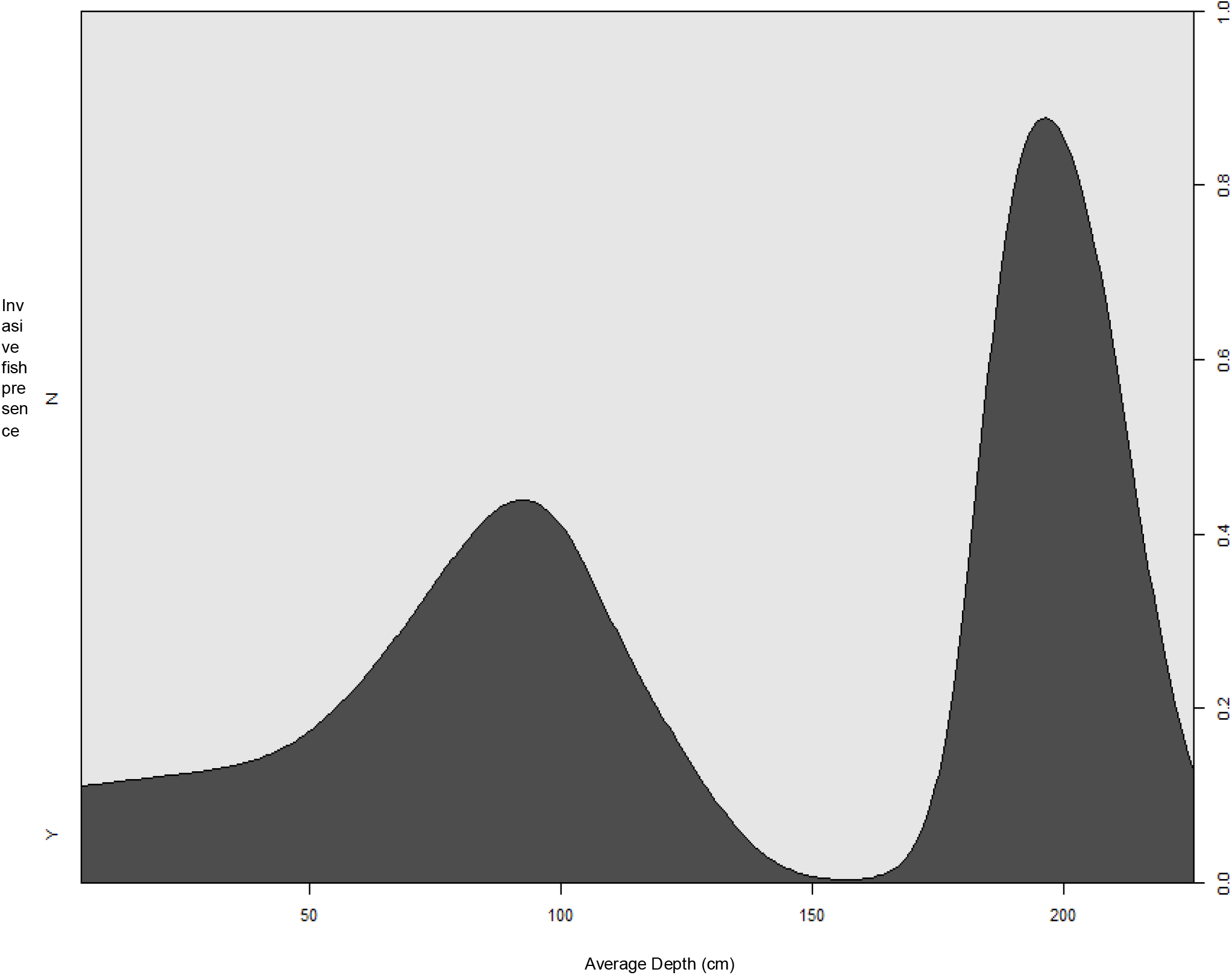
Critical Difference Plot illustrating the proportion of the binomial outcome for invasive fish presence (Yes= Dark Grey, No=Light Grey) against the average depth (cm) of sites sampled *for ghost frog tadpoles in the southern Cape Fold Mountains*.

## Literature

Akinwande MO, Dikko HG, Samson A. 2015. Variance Inflation Factor: As a Condition for the Inclusion of Suppressor Variable(s) in Regression Analysis. Open Journal of Statistics 05:754–767. DOI: 10.4236/ojs.2015.57075.

Alford RA, Richards SJ. 1999. Global Amphibian Declines: A Problem in Applied Ecology. Annual Review of Ecology and Systematics 30:133–165. DOI: 10.1146/annurev.ecolsys.30.1.133.

Angus O, Turner AA, Measey J. 2023. In a Rough Spot: Declines in Arthroleptella rugosa calling densities are explained by invasive pine trees. Austral Ecology. DOI: 10.1111/aec.13273.

Avidon S, Shelton JM, Marr SM, Bellingan TA, Esler KJ, Weyl OLF. 2018. Preliminary evaluation of non-native rainbow trout (Oncorhynchus mykiss) impact on the Cederberg ghost frog (Heleophryne depressa) in South Africa’s Cape Fold Ecoregion. African Journal of Aquatic Science 43:313–318. DOI: 10.2989/16085914.2018.1507898.

Bellingan TA, Hugo S, Woodford DJ, Gouws J, Villet MH, Weyl OLF. 2019. Rapid recovery of macroinvertebrates in a South African stream treated with rotenone. Hydrobiologia 834:1–11. DOI: 10.1007/s10750-019-3885-z.

Brooks ME, Kristensen K, Benthem KJ, Magnusson A, Berg CW, Nielsen A, Skaug HJ, Mächler M, Bolker BM. 2017. glmmTMB Balances Speed and Flexibility Among Packages for Zero-inflated Generalized Linear Mixed Modeling. The R Journal 9:378. DOI: 10.32614/RJ-2017-066.

Bucciarelli GM, Blaustein AR, Garcia TS, Kats LB. 2014. Invasion Complexities: The Diverse Impacts of Nonnative Species on Amphibians. Copeia 2014:611–632. DOI: 10.1643/OT-14-014.

Cambray JA. 2003. The global impact of alien trout species — a reviewlll; with reference to their impact in South Africa The global impact of alien trout species — a reviewlll; with reference to their. African Journal of Aquatic Science 28:61–67. DOI: 10.2989/16085914.2003.9626601.

Chakona A, Swartz ER, Gouws G. 2013. Evolutionary Drivers of Diversification and Distribution of a Southern Temperate Stream Fish Assemblage: Testing the Role of Historical Isolation and Spatial Range Expansion. PLOS ONE 8:e70953. DOI: 10.1371/journal.pone.0070953.

Clark KL, Lazerte BD. 1985. A Laboratory Study of the Effects of Aluminum and pH on Amphibian Eggs and Tadpoles. Canadian Journal of Fisheries and Aquatic Sciences 42:1544–1551. DOI: 10.1139/f85-193.

Collins JP. 2010. Amphibian decline and extinction: What we know and what we need to learn. Diseases of Aquatic Organisms 92:93–99. DOI: 10.3354/dao02307.

Collins JP, Storfer A. 2003. Global amphibian declines: sorting the hypotheses. Diversity & Distributions 9:89–98. DOI: 10.1046/j.1472-4642.2003.00012.x.

Conradie W, Conradie C. 2015. Correlation between development and increase of number of labial tooth rows in Ghost Frog tadpoles (Anura: Heleophrynidae). Acta Herpetologica 10:143–148. DOI: 10.13128/Acta_Herpetol-16427.

Cox JG, Lima SL. 2006. Naiveté and an aquatic–terrestrial dichotomy in the effects of introduced predators. Trends in Ecology & Evolution 21:674–680. DOI: 10.1016/j.tree.2006.07.011.

Craney TA, Surles JG. 2002. Model-dependent variance inflation factor cutoff values. Quality Engineering 14:391–403. DOI: 10.1081/QEN-120001878.

Denoel M, Dzukic G, Kalezic ML. 2005. Effects of Widespread Fish Introductions on Paedomorphic Newts in Europe. Conservation Biology 19:162–170. DOI: 10.1111/j.1523-1739.2005.00001.x.

Ebrahim Z, de Villiers A, Measey J. 2020. Assessing water conditions for heleophryne rosei tadpoles and the conservation relevance. Koedoe 62:1–6. DOI: 10.4102/KOEDOE.V62I1.1581.

Ellender BR. 2013. Ecological consequences of non-native fish invasion in eastern Cape headwater streams. PhD Thesis. Rhodes University.

Ellender BR, Wasserman RJ, Chakona A, Skelton PH, Weyl OLF. 2017. A review of the biology and status of Cape Fold Ecoregion freshwater fishes. Aquatic Conservation: Marine and Freshwater Ecosystems 27:867–879. DOI: 10.1002/aqc.2730.

Ellender BR, Weyl OLF. 2014. A review of current knowledge, risk and ecological impacts associated with non-native freshwater fish introductions in South Africa. Aquatic Invasions 9:117–132. DOI: 10.3391/ai.2014.9.2.01.

Falaschi M, Giachello S, Lo Parrino E, Muraro M, Manenti R, Ficetola GF. 2021. Long-term drivers of persistence and colonization dynamics in spatially structured amphibian populations. Conservation Biology 35:1530–1539. DOI: 10.1111/cobi.13686.

Falaschi M, Melotto A, Manenti R, Ficetola GF. 2020. Invasive Species and Amphibian Conservation. Herpetologica 76:216–227. DOI: 10.1655/0018-0831-76.2.216.

Fox J, Weisberg S. 2019. An R companion to applied regression. Thousand Oaks CA: Sage.

Gillespie GR. 2001. The role of introduced trout in the decline of the spotted tree frog (Litoria Spenceri) in south-eastern Australia. Biological Conservation 100:187–198. DOI: 10.1016/S0006-3207(01)00021-0.

Green DM, Lannoo MJ, Lesbarrères D, Muths E. 2020. Amphibian population declines: 30 years of progress in confronting a complex problem. Herpetologica 76:97–100. DOI: 10.1655/0018-0831-76.2.97.

Green M, Thompson MB, Lemckert FL. 2004. The effects of suspended sediments on the tadpoles of two stream-breeding and forest dwelling frogs,Mixophyes balbus andHeleioporus australiacus

Karssing RJ, Rivers-moore NA, Slater K. 2012. Influence of waterfalls on patterns of association between trout and Natal cascade frog Hadromophryne natalensis tadpoles in two headwater streams in the uKhahlamba Drakensberg Park World Heritage Site, South Africa. African Journal of Aquatic Science 5914:107–112. DOI: 10.2989/16085914.2012.666381.

Karssing R, Rivers-Moore N, Slater K. 2012. Influence of waterfalls on patterns of association between trout and Natal cascade frog Hadromophryne natalensis tadpoles in two headwater streams in the uKhahlamba Drakensberg Park World Heritage Site, South Africa. African Journal of Aquatic Science 37:107–112. DOI: 10.2989/16085914.2012.666381.

Kats LB, Ferrer RP. 2003. Alien predators and amphibian declines: Review of two decades of science and the transition to conservation. Diversity and Distributions 9:99–110. DOI: 10.1046/j.1472-4642.2003.00013.x.

Khosa D, Marr SM, Wasserman RJ, Zengeya TA, Weyl OLF. 2019. An evaluation of the current extent and potential spread of Black Bass invasions in South Africa. Biological Invasions 21:1721–1736. DOI: 10.1007/s10530-019-01930-0.

Loppnow G, Vascotto K, Venturelli P. 2013. Invasive smallmouth bass (Micropterus dolomieu): history, impacts, and control. Management of Biological Invasions 4:191–206. DOI: 10.3391/mbi.2013.4.3.02.

Lukas P. 2021. Larval cranial anatomy of the Eastern Ghost Frog (Heleophryne orientalis). Acta Zoologica 102:452–466. DOI: 10.1111/azo.12352.

Martín-Torrijos L, Sandoval-Sierra JV, Muñoz J, Diéguez-Uribeondo J, Bosch J, Guayasamin JM. 2016. Rainbow trout (Oncorhynchus mykiss) threaten Andean amphibians. Neotropical Biodiversity 2:26– 36. DOI: 10.1080/23766808.2016.1151133.

Measey GJ. 2011. Ensuring a future for south africa’s frogs: a strategy for conservation research. In: Measey GJ ed. SANBI biodiversity series 19. Pretoria: South African National Biodiversity Institute, 2–6. DOI: 10.1080/0035919x.2011.586444.

Measey J, Tarrant J, Rebelo A, Turner A, Du Preez L, Mokhatla M, Conradie W. 2019. Has strategic planning made a difference to amphibian conservation research in South Africa? Bothalia 49. DOI: 10.4102/abc.v49i1.2428.

Melotto A, Ficetola GF, Alari E, Romagnoli S, Manenti R. 2021. Visual recognition and coevolutionary history drive responses of amphibians to an invasive predator. Behavioral Ecology 32:1352–1362.

Midgley J, Schafer G. 1992. Correlates of water colour in streams rising in Southern Cape catchments vegetated by fynbos and/or forest. Water SA 18:93–100.

Minter LR, Burger M, Harrison JA, Braak HH, Bishop PJ, Kloepfer D (eds.). 2004. Atlas and red data book of the frogs of South Africa, Lesotho, and Swaziland. Washington DC: Smithsonian Institution.

Miró A, Sabás I, Ventura M. 2018. Large negative effect of non-native trout and minnows on Pyrenean lake amphibians. Biological Conservation 218:144–153. DOI: 10.1016/j.biocon.2017.12.030.

Moore DR, Spittlehouse DL, Story A. 2005. Riparian Microclimate and Stream Temperature Response to Forest Harvesting: A Review1. JAWRA Journal of the American Water Resources Association 41:813–834. DOI: 10.1111/j.1752-1688.2005.tb03772.x.

Mucina L, Rutherford MC (eds.). 2006. The vegetation of South Africa, Lesotho and Swaziland. Pretoria: South African National Biodiversity Institute.

Nunes AL, Fill JM, Davies SJ, Louw M, Rebelo AD, Thorp CJ, Vimercati G, Measey J. 2019. A global meta-analysis of the ecological impacts of alien species on native amphibians. Proceedings of the Royal Society B: Biological Sciences 286.

R Core Team. 2021. R: A language and environment for statistical computing.

Remon J, Bower DS, Gaston TF, Clulow J, Mahony MJ. 2016. Stable isotope analyses reveal predation on amphibians by a globally invasive fish (Gambusia holbrooki). Aquatic Conservation: Marine and Freshwater Ecosystems 26:724–735. DOI: 10.1002/aqc.2631.

Richardson DM, Holmes PM, Esler KJ, Galatowitsch SM, Stromberg JC, Kirkman SP, Pyšek P, Hobbs RJ. 2007. Riparian vegetation: degradation, alien plant invasions, and restoration prospects: Riparian vegetation: degraded, invaded, transformed. Diversity and Distributions 13:126–139. DOI: 10.1111/j.1366-9516.2006.00314.x.

Rivers-Moore NA, Fowles B, Karssing RJ. 2013. Impacts of trout on aquatic macroinvertebrates in three Drakensberg rivers in KwaZulu-Natal, South Africa. African Journal of Aquatic Science 38:93–99. DOI: 10.2989/16085914.2012.750592.

Rivers-Moore NA, Karssing RJ. 2014. Water temperature affects life-cycle duration of tadpoles of Natal cascade frog. African Journal of Aquatic Science 39:223–227. DOI: 10.2989/16085914.2014.903165.

Rowntree K, Wadeson R. 2000. Field manual for channel classification and condition assessment. Institute for Water Quality Studies, Department of Water Affairs and Forestry, Pretoria, South Africa.

SAFRoG SAFRG (SA-Fr (IUCN, IUCN-ASG ISASG (IUCN. 2016a. IUCN Red List of Threatened Species: Heleophryne hewitti. IUCN Red List of Threatened Species.

SAFRoG SAFRG (SA-Fr (IUCN, IUCN-ASG ISASG (IUCN. 2016b. IUCN Red List of Threatened Species: Heleophryne rosei. IUCN Red List of Threatened Species.

Scott DF, Prinsloo FW. 2008. Longer-term effects of pine and eucalypt plantations on streamflow. Water Resources Research 44:1–8. DOI: 10.1029/2007WR006781.

Seebens H, Blackburn TM, Dyer EE, Genovesi P, Hulme PE, Jeschke JM, Pagad S, Pyšek P, Winter M, Arianoutsou M, Bacher S, Blasius B, Brundu G, Capinha C, Celesti-Grapow L, Dawson W, Dullinger S, Fuentes N, Jäger H, Kartesz J, Kenis M, Kreft H, Kühn I, Lenzner B, Liebhold A, Mosena A, Moser D, Nishino M, Pearman D, Pergl J, Rabitsch W, Rojas-Sandoval J, Roques A, Rorke S, Rossinelli S, Roy HE, Scalera R, Schindler S, Štajerová K, Tokarska-Guzik B, van Kleunen M, Walker K, Weigelt P, Yamanaka T, Essl F. 2017. No saturation in the accumulation of alien species worldwide. Nature Communications 8:14435. DOI: 10.1038/ncomms14435.

Shaffer TL, Johnson DH. 2008. Ways of Learning: Observational Studies Versus Experiments. The Journal of Wildlife Management 72:4–13. DOI: 10.2193/2007-293.

Shelton JM, Samways MJ, Day JA. 2015. Predatory impact of non-native rainbow trout on endemic fish populations in headwater streams in the Cape Flortistic Region of South Africa. Biological Invasions 17:365–379. DOI: 10.1007/s10530-014-0735-9.

Sutherland WJ, Pullin AS, Dolman PM, Knight TM. 2004. The need for evidence-based conservation. Trends in Ecology & Evolution 19:305–308. DOI: 10.1016/j.tree.2004.03.018.

Velasco MA, Berkunsky I, Simoy M V., Quiroga S, Bucciarelli G, Kats L, Kacoliris FP. 2018. The rainbow trout is affecting the occupancy of native amphibians in Patagonia. Hydrobiologia 817:447–455. DOI: 10.1007/s10750-017-3450-6.

Wake DB, Vredenburg VT. 2008. Are we in the midst of the sixth mass extinction? A view from the world of amphibians. Proceedings of the National Academy of Sciences 105:11466–11473. DOI: https://doi.org/10.1073/pnas.0801921105.

van der Walt JA, Marr SM, Wheeler MJ, Impson DN, Garrow C, Weyl OLF. 2019. Successful mechanical eradication of spotted bass (Micropterus punctulatus (Rafinesque, 1819)) from a South African river. Aquatic Conservation: Marine and Freshwater Ecosystems f:1–9. DOI: 10.1002/aqc.3035.

Wassersug R. 2000. Tadpoles: The Biology of Anuran Larvae. Copeia 2000:1125–1134. DOI: 10.1643/0045-8511(2000)000[1125:BR]2.0.CO;2.

Weyl OLF, Finlayson B, Impson ND, Woodford DJ, Steinkjer J. 2014a. Threatened Endemic Fishes in South Africa’s Cape Floristic Region: A New Beginning for the Rondegat River. Fisheries 39:270–279. DOI: 10.1080/03632415.2014.914924.

Weyl OLF, Finlayson B, Impson ND, Woodford DJ, Steinkjer J. 2014b. Threatened endemic fishes in south africa’s cape floristic region: A new beginning for the rondegat river. Fisheries 39:270–279. DOI: 10.1080/03632415.2014.914924.

Weyl PSR, de Moor FC, Hill MP, Weyl OLF. 2010. The effect of largemouth bass Micropterus salmoides on aquatic macroinvertebrate communities in the Wit River, Eastern Cape, South Africa. African Journal of Aquatic Science 35:273–281. DOI: 10.2989/16085914.2010.540776.

Woodford DJ, Impson ND. 2004. A preliminary assessment of the impact of alien rainbow trout (Oncorhynchus mykiss) on indigenous fishes of the upper Berg River, Western Cape Province, South Africa. African Journal of Aquatic Science 29:107–111. DOI: 10.2989/16085910409503799.

Woodford DJ, Impson ND, Day JA, Bills IR. 2005a. The predatory impact of invasive alien smallmouth bass, Micropterus dolomieu (Teleostei: Centrarchidae), on indigenous fishes in a Cape Floristic Region mountain stream. African Journal of Aquatic Science 30:167–173. DOI: 10.2989/16085910509503852.

Woodford DJ, Impson ND, Day JA, Bills IR. 2005b. The predatory impact of invasive alien smallmouth bass, Micropterus dolomieu (Teleostei: Centrarchidae), on indigenous fishes in a Cape Floristic Region mountain stream. African Journal of Aquatic Science 30:167–173. DOI: 10.2989/16085910509503852.

Zuur AF, Ieno EN, Elphick CS. 2010. A protocol for data exploration to avoid common statistical problems. Methods in Ecology and Evolution 1:3–14. DOI: 10.1111/j.2041-210X.2009.00001.x.

